# Single-cell histories in growing populations: relating physiological variability to population growth

**DOI:** 10.1101/100495

**Authors:** Philipp Thomas

## Abstract

Cell size and individual growth rates vary substantially across genetically identical cell populations. This variation cannot entirely be explained by asynchronous cell division cycles, but also needs to take into account the differences in the histories that cells experience during their lifespan. We describe a stochastic framework to characterise cell size histories in an exponentially growing population. We show that these histories differ from cells observed in isolation, such as observed in mother machines. Quantifying these historical fluctuations allows us to predict the population growth rate. We highlight that the maximum attainable population growth cannot exceed the rate at which an average cell grows, but the population doubles faster than an average cell doubles its size. We validate this prediction using recent single-cell data. The theory thus provides fundamental limits on population fitness in terms of individual cell properties.

## I. INTRODUCTION

Clonal populations often exhibit a high degree of variability in cellular physiology^1–4^. Analysing the sources of these variations remains challenging because the statistics of single cells observed in isolation often differ from those observed in growing and dividing populations. In the absence of environmental and genetic variation, two factors determine phenotypic variability: (i) sister cells differ in cell cycle duration, growth rate and cell size because the intracellular processes by which these are determined operate ultimately using finite number of molecules^5,6^, while (ii) distantly related cells differ because they experienced different life histories^7^. Dissecting these effects is crucial to infer what fluctuations cells are facing during their replicative lifespans^8^.

Variation between sister cells is commonly studied by acquiring lineage trajectories that follow either one of the sister cells in each division while discarding the information about the other sister. We will refer to such a lineage as a *forward lineage* shown in Fig. I (blue). Experimentally, such observations are obtained using microfluidic traps^9,10^ or using computational cell tracking^5^, which have now been extensively used to inform phenomenological models for the dynamics of growth and division cycles of individual cells^2,11,12^. The benefit of this approach is that the age of each cell can be determined accurately without the bias that is observed in populations due to distributed cell ages and sizes.

**FIG. 1.**
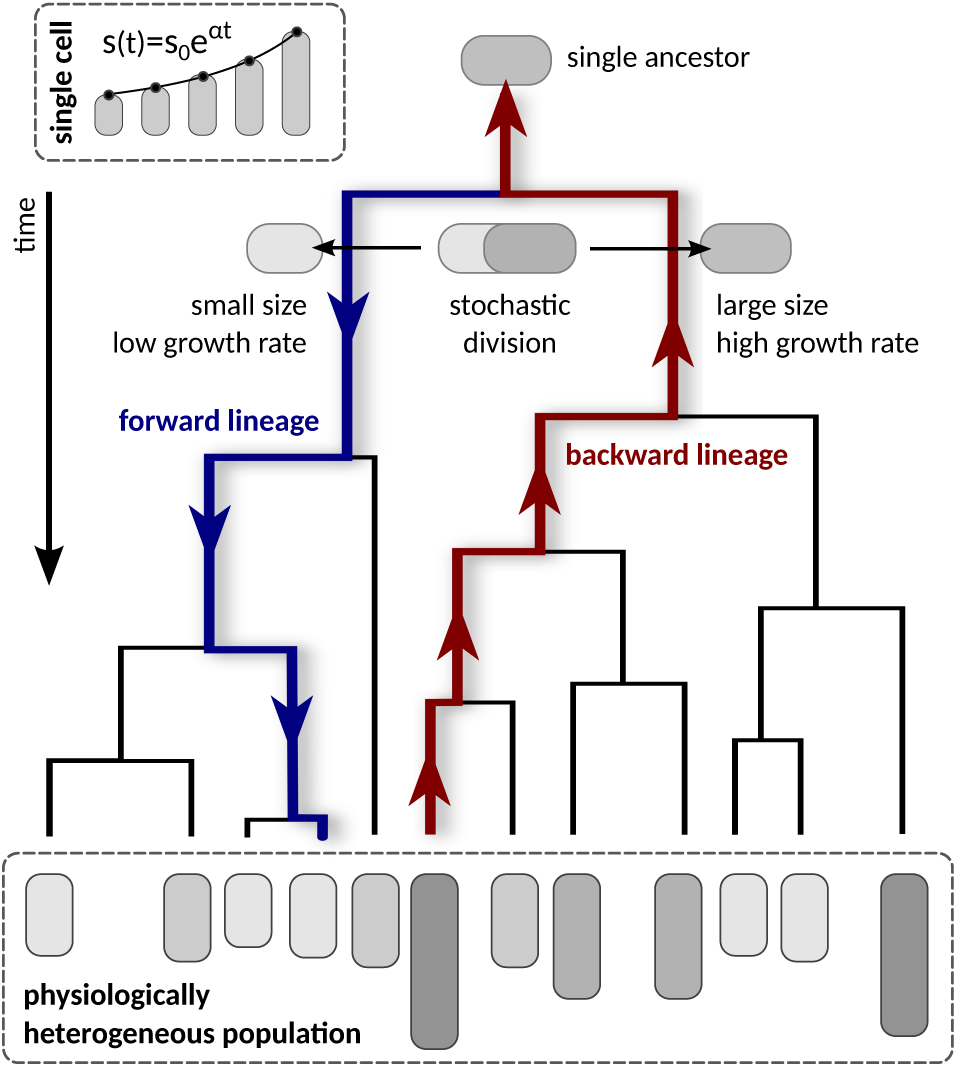
Lineages of clonal populations. A population with distributed cell sizes arises from a pedigree with a common ancestor. Two types of lineages are used to characterise the tree. Forward lineages start from a common ancestor, end at an arbitrary cell in the population, and hence correspond to possible cell fates. Backward lineages start at an arbitrarily chosen cell in the population, end at a common ancestor, and are representative of a cell’s history.

Observing the variation in the histories of single cells is more elaborate because one needs to track an ensemble of growing and dividing cells to construct lineage trees of whole populations^1,8,13,14^. In this way, one resolves the life histories of all individuals in the population preserving their ancestral relationships. To be precise, we here consider a cell’s history which represents a lineage obtained from choosing an arbitrary cell in a population and tracing it back to the common ancestor the population originated from, see Fig. I (red line). For each cell in the population, there exists exactly one such path, which we will henceforth denote as a *backward lineage*. It is to be expected that the distribution of these trajectories represent a typical cell’s history.

While most forward lineages may correspond to a possible cell history, it is unclear whether a typical history can be predicted from a forward lineage, because it ignores ancestral relationships. Intuitively one might argue that each history contributes differentially to the overall population growth. A typical history may diverge from a lineage of a single cell observed in isolation even when environmental stresses or limitations are absent. Quantifying this dependence can give valuable insights into how single-cell parameters shape individual cell histories, how these properties are represented in a clonal population (Fig. I dashed box), and thus how cells keep memories of their past^15^.

Theoretical studies have mainly focussed on the forward approach because it can be simulated without affording to track whole populations^11^ and because it is often more amenable to analysis^16–19^. Retrospective models of populations are more common when studying genetic diversity due to rare mutations in populations with age-structure^20,21^. Few theoretical studies to-date have given a quantitative description of cell histories due to phenotypic fluctuations^22,23^. These studies established that a typical cell’s history determines the population-growth rate but also that the strength of selection acting on an individual can be inferred from this history^8,23^. Such an analysis is limited to the study of individual differences in cell age, which makes it difficult to reason about variation in cellular physiology that is typically observed in clonal populations.

While cellular division times are highly variable, division timing is not entirely determined by cell age, but also by cell growth and size. Cell division in bacteria, for instance, follows after elongation by a stochastic amount of cell length^11,17,24^, while other microbes more closely resemble sizer or timer controls^25^. The fact that each cell inherits a certain fraction of the mother’s cell size, leading to correlated division times, complicates analysis of these effects. It, therefore, remains an open question what is the effect of stochasticity in cell physiology on individual cell histories in a growing population, and what part of this cell-to-cell variability contributes to the population-growth rate that is commonly associated with the *fitness* of a population.

In this article, we compare the lineage statistics obtained from an exponentially growing population whose individuals control division timings based on cell size. First, we briefly introduce our model assumptions and discuss the concepts of forward and backward lineages. We show that the population-growth rate is constrained by the distributions of birth-size and growth histories while relegating technical details to the Methods section. We then use these result to study how differences between sister cells affect typical cell histories. Specifically, in Sec. IIA we study a population of cells whose individuals divide into two perfectly identical sister cells inheriting equal proportions of the mother’s size. For these cases, the statistics of forward and backward lineages agree but it is not possible to maintain a stable size distribution despite the stochasticity in division times. In Sec. IIB we investigate the consequences of imperfect division due to division errors. The cell-growth rate limits population-growth rate and is independent of the size control. The implications for asymmetrically dividing cell populations and the contributions of individuals to the overall population growth are discussed. Finally, in Sec. IIC we investigate the effects of sister cell variations in the cell-growth rate. Using individual cell histories we show that growth rate fluctuations are subject to negative selection and provide bounds on the maximum attainable population-growth. We also investigate the effect of asymmetric sister cell growth. We finish with a discussion where we elaborate on applications of our findings in the context of recent single cell techniques. Our analysis highlights that cell histories in a population are strongly affected by differences between sister cells arising from the cell division process.

## II. RESULTS

We assume that each individual cell undergoes exponential growth, a dependence that has been reported for *E. coli*^9^, *C. crescentus*^24^, *B. subtilis*^2^ and *S. cerevisiae*^12^.

The size of a cell of age *τ* follows the deterministic relation

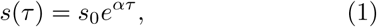

where *s*_0_ is the size at birth and *α* is the exponential cell-growth rate. We account for three sources of variability: fluctuations in the birth size *s*_0_ and division size *s_d_*, as well as variations in the cell-growth rate *α* between division cycles. Such growth variations have been observed in bacteria whose increase in length follows a single exponential between birth and division^2,4,9,26^.

Assuming that cell division occurs at a rate *γ*(*τ*, *s*,*α*), it follows that the division time *τ_d_* is distributed according to

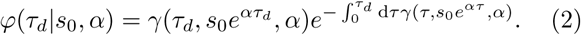

Changing variables from division time *τ_d_* to division size *s_d_* using Eq. (1), we find

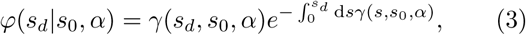

where we set *γ*(*τ_d_*, *s_d_*, *α*) = *αsγ*(*s_d_*, *s*_0_, *α*). Stochasticity in the division times can therefore be directly related to fluctuations in the division size, and vice versa. The division size representation is convenient because it can be used to model cell size control, which has been reported to vary with growth media but is relatively independent of cell-growth rate^2^. We adopt this independence assumption from here on.

### Forward lineages

The lineage approach is agnostic about population growth because it follows only of the two daughter cells. It is therefore not difficult to write an equation for the distribution of birth sizes and growth rates in a forward lineage^16,19^ (Methods 1), *ψ_fw_*(*s*_0_,*α*), which is

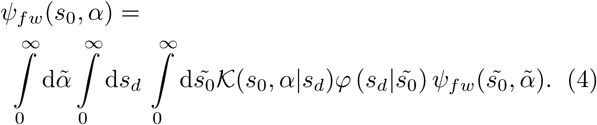

The division kernel *κ*(*s*_0_, *α*|*s_d_*) is the distribution of birth size and growth rate of the progenies of a mother cell with size *s_d_*. When division produces two daughters the it is given by a mixture

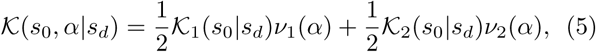

where *κ*_1,2_(*s*_0_|*s_d_*) denotes the distribution of birth size after cell division and *ν*_1,2_(*α*) the distribution of growth rates, which we assume to be independent^2,27^. We do not consider inheritance of growth rates because experiments in bacteria showed that correlations in growth rates are negligible across generations^2,4,9^.

### Cell histories

Determining the statistics of cell histories is more elaborate because we need to account for the growth of the whole cell population. To this end we consider the density *n*(*τ, s, α, t*) that counts the number of cells with age *τ*, size *s*, and cell-growth rate *α*. The total number of cells is given by

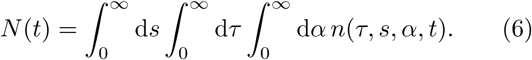

Since the probability for a cell to divide at age *τ* and size *s* is given by *γ*(*τ, s, α*)d*τ*d*s*d*α*, the cell density follows the evolution equation

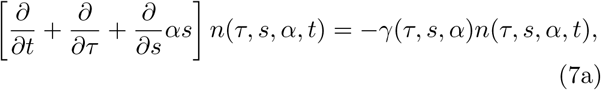

Assuming further that the cell-growth rate *α* changes only at cell division, the density obeys the boundary condition

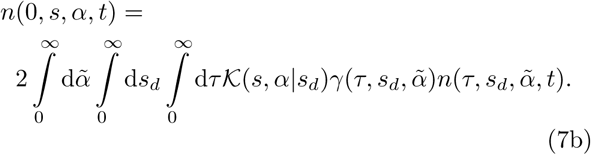

The condition ensures that the number of newborn cells in the population must equal twice the number of dividing cells. Similar approaches have been used to characterise distributions of age and size-structured populations^28–34^.

Eqs. (7) are difficult to solve in practise because of the simultaneous dependence of the division rate on age and size. To make analytical progress, we restrict ourselves to the long-term behaviour in which the total cell number grows exponentially

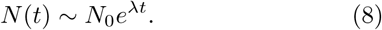

Also the total mass of the population grows at the same rate. We show (Methods 2) that the distribution of birth size and cell-growth rates in backward lineages is obtained from the solution of Eqs. (7) as follows

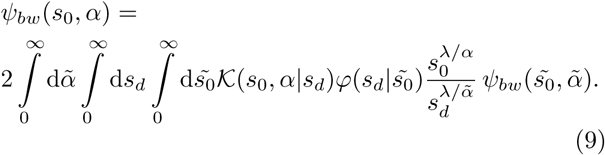

Solving for the backward lineage distribution *ψ*_*bw*_ allows us to predict the distributions measured across a growing population (Methods 2, Eq. (37a)).

The population-growth rate λ, however, is a priori unknown. It can be shown that it is given by the maximum value of λ for which *ψ*_*bw*_ can be normalised (Methods 2), and hence the backward lineage contains the full information about the population growth. We note that if there is a unique λ with dominant real part, the fraction of cells with a certain age and size is constant in a growing population, i.e. it settles to a stable size distribution. We will show how solutions to this equation are obtained in biologically relevant cases. We begin by studying stochasticity in the cell size control (Sec. II A), and then include variations in inherited cell size (Sec. II B) and cell-growth rates (Sec. II C).

### A. Populations of perfectly dividing cells

As a first example we investigate an idealised situation in which each mother cell divides exactly into two identical daughter cells, which entails that both sizes and both cell-growth rates must be identical for the two daughter cells. This shows that albeit stochasticity in size control may lead to a stable age distribution, it cannot achieve a stable size distribution in a population.

#### No stable size distribution for perfectly dividing cells

Letting *κ*(*s*_0_, *α*|*s_d_*) = *δ*(*s*_0_ – *s_d_*/2)*δ*(*α* –*α*_0_) in Eq. (9) reduces the equation to the integral

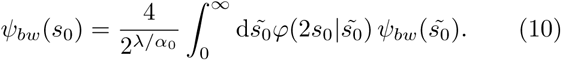

We obtain the characteristic equation for λ by integrating over *s*_0_ and using the fact that *φ* is normalised. This leads to 2 = 2^λ/*α*0^, which has the roots 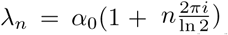 for *n* = 0,1,…. Since there exists no single value with dominant real part, the size distribution of perfectly dividing cells oscillates indefinitely. While this result is well known for cells that divide at a critical size^35^, we find that it holds for arbitrary cell size controls.

#### Ideal cell histories

Although a stable size distribution does not exist, we investigate the distribution corresponding to the population
growing at rate λ = *α*. This distribution can be considered as the limiting backward lineage distribution for small but non-zero division errors or cell-growth rate variability, which satisfies

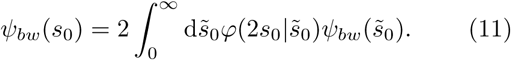

The equation is same as Eq. (4) for *ψ_fw_*, and therefore cell histories and forward lineages are identically distributed. The integral can be carried out if *φ* is independent of 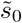 for which *ψ_bw_*(*s*_0_) = 2*φ*(2*s*_0_). However, more generally the lineage distribution cannot be obtained without knowing the explicit form of the size control *φ*.

### B. Populations of imperfectly dividing cells

Although cell division is a tightly controlled process it is hard to imagine that a mother cell can divide into two identically sized daughters. Even in symmetrically dividing organisms as *E. coli* the inherited size differs between daughter cells^36–38^. In this situation, the division kernel *κ*(*s*_0_,*α*|*s_d_*) in Eq. (5) can be written as

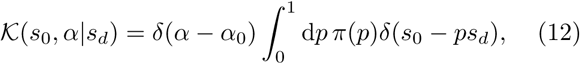

where *p* denotes the ratio of daughter to mother cell size.

#### Cell-growth rate sets population-growth rate

To characterise the population growth, we derive a characteristic equation for λ. Inserting Eq. (12) into Eq. (9) and integrating over *s*_0_ yields

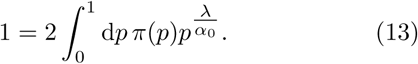

The inherited size fraction obeys *π*(*p*) = *π*(1 − *p*) because of size conservation. This implies that the daughter cells inherited half the mother size on average, i.e. 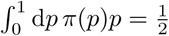, and thus the real solution to this equation is λ = *α*_0_. For cell populations producing daughters of different sizes, it follows that the elongation rate alone determines the population-growth rate.

#### Cells are bigger in backward lineages

Next, we study the distribution of birth sizes in backward lineages. Using the result of the previous paragraph together with Eq. (12) we find

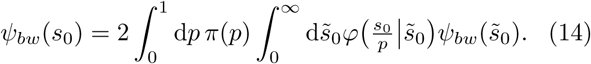

We note that this distribution is different than the one of a forward lineage. Specifically, comparison with Eq. (4) shows that the ratio of daughter to mother size in a backward lineage is distributed as

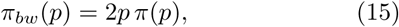

rather than *π*(*p*) as in the forward lineage. The equation shows that the observed daughter-mother ratio is skewed towards bigger daughters. Intuitively, this bias can be understood by the fact that bigger cells grow faster, divide quicker, and thus are overrepresented in backward lineages.

To investigate this effect using stochastic simulations we use a linear model relating division size to the size at birth of single cells. Many experimental studies employed such a linear regression to quantify division control^11,25^. To this end we set *γ*(*s*, *s*_0_) = *γ*(*s* − *as*_0_), or equivalently

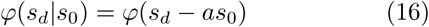

from which the division sizes can be sampled. The parameter *a* denotes different models of cell size control. For *a* = 0 division size varies irrespectively of birth size, often called the sizer mechanism. For *a* = 1 the size added from birth to division varies independently of birth size, an adder mechanism that is commonly observed in bacteria^2,17,24^. For *a* = 2 division size is proportional to birth size, which resembles cell-size control based on division timing. Intermediate values would behave either sizer- or timer-like^2^. Using the latter dependence we study the case of large division errors corresponding to an almost uniform distribution *π*(*p*) shown in Fig. 2a (solid black line). We can compare this distribution to *π_bw_*(*p*) which yields the daughter-mother ratio in backward lineages (solid red line). The stochastic simulations (shaded blue area) are in excellent agreement with the theoretical predictions.

**FIG. 2.**
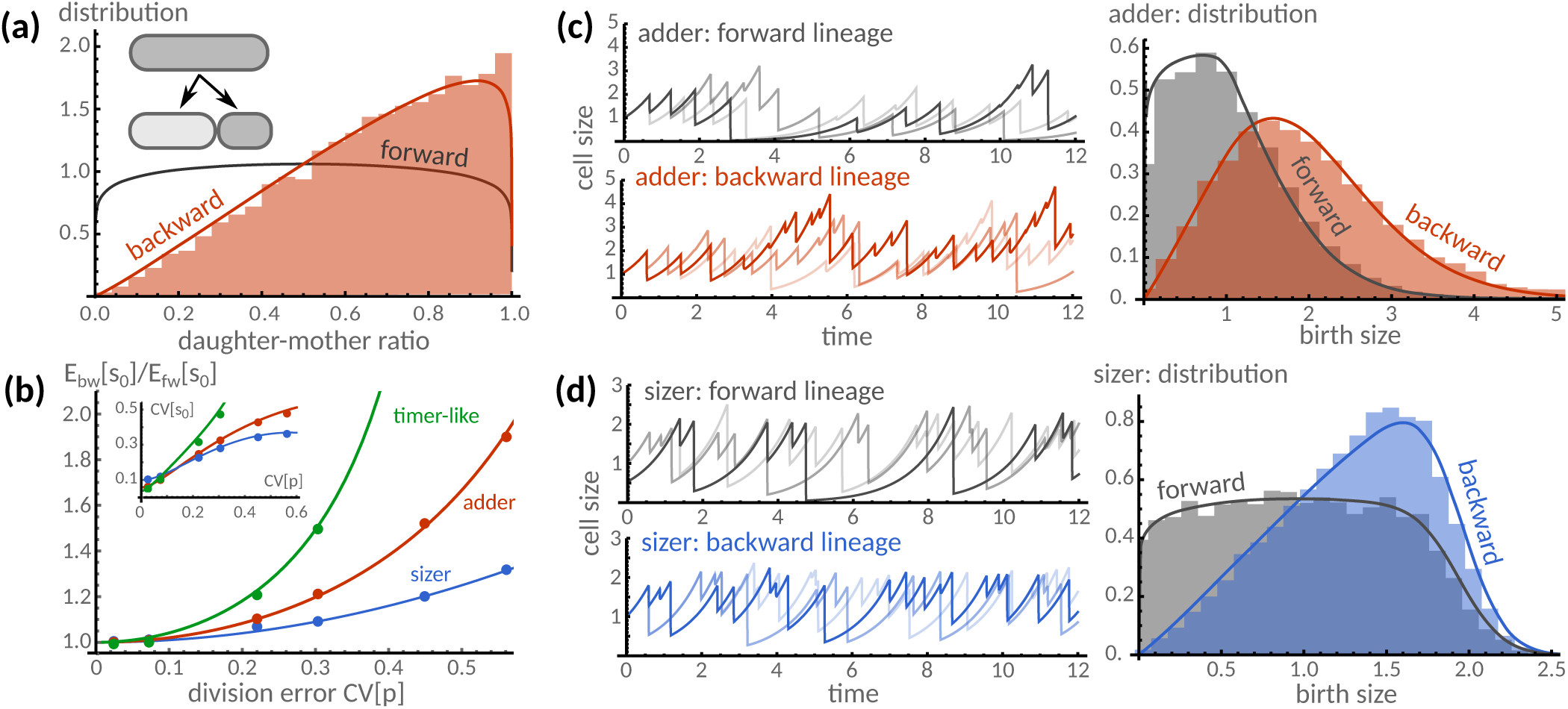
Division errors increase cell size in backward lineages. **(a)** The distribution of daughter to mother-size ratios is compared for the two lineage types. Because each mother produces two offspring, the distribution of the forward lineage is symmetric. In the backward lineages the same quantity is significantly skewed towards larger daughter cells. **(b)** The average birth size increases with increasing division error CV[*p*]. This amplification is smaller for the sizer model than for adder and timer-like models (see main text for model details). The coefficient of variation is shown in the inset. **(c)** We compare forward and backward lineage traces (left) collected from a clonal population implementing the adder size control. We observe that cells are larger in backward than in forward lineages. We quantify this dependence using the distributions of birth sizes in these lineages (right) obtained from theoretical predictions, numerical solution of Eq. (14) (solid red line) and Eq. (4) (solid black), and stochastic simulations (shaded areas). **(d)** For the sizer mechanism (left) both lineage types show comparable ranges of cell sizes but cells in backward lineages divided more frequently and at larger sizes. Our theoretical prediction (right) confirms this dependence showing that larger cells are more frequently observed in backward lineages than in forward ones. Theory and simulations assumed division errors to follow a symmetric Beta distribution with a coefficient of variation of about 66% while the error in size control was modelled using a Gamma distribution with coefficient of variation of 10%.

A general solution of Eq. (14) is, however, difficult to obtain. We therefore characterise the distribution by its moments. The first moment, for example, can be obtained exactly for linear models, Eq. (16), through multiplying Eq. (14) by *s*_0_, integrating and solving the expression for the expectation value of *s*_0_. The mean birth size is obtained as

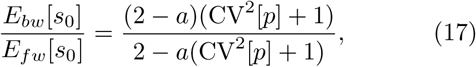

where CV[*p*] denotes the “division error” given by the coefficient of variation of *π*(*p*). Interestingly, we find that, while in forward lineages, the mean birth size is independent of the division error, birth size increases with division errors in backward lineages. When these errors are small, the increase in birth size is solely determined by the division error and independent of the mechanism of size control. For higher division errors, we find that birth size increases with *a* (Fig. 2b). Hence an adder mechanism leads to larger cells than the sizer mechanism. Likewise a timer-like mechanism (*a* = 1.5) produces larger cells than the adder control.

To demonstrate this counter-intuitive phenomenon in more detail, we performed stochastic simulations. Three representative lineages, shown for the adder (c, left) and sizer mechanisms (d, left), demonstrate that cells observed in backward lineages are typically bigger than in forward lineages. The histograms of birth sizes, obtained from simulations and by numerically solving Eq. (14), confirm that cells are generally bigger in backward lineages than in forward lineages, while the distribution obtained using the adder (c, right) is significantly wider than the one obtained using the sizer control (d, right).

#### Histories of asymmetrically dividing cells have more mother than daughters

Albeit two daughter cells are never truly identical, for many species cell division is naturally asymmetric. In budding yeast, for instance, cell division distinguishes a smaller daughter from a mother cell. In this paragraph, we refer to the bigger cell as a mother, and to the smaller cell as a daughter and specify this dependence in the division size distributions. In forward lineages, mother (m) and daughter cells (d) can only occur in equal proportions

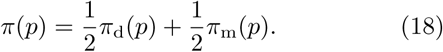

Because total size is conserved in the division process, these distributions obey *π*_m_(*p*) = *π*_d_(1 − *p*). The distribution of division size ratios is bimodal (Fig. 3a, dashed grey line). From the definition of *π*_*bw*_, Eq. (15), it follows that the proportion of size inherited by either cell type determines the fraction of mother cells observed in backward lineages as follows

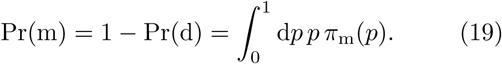

**FIG. 3.**
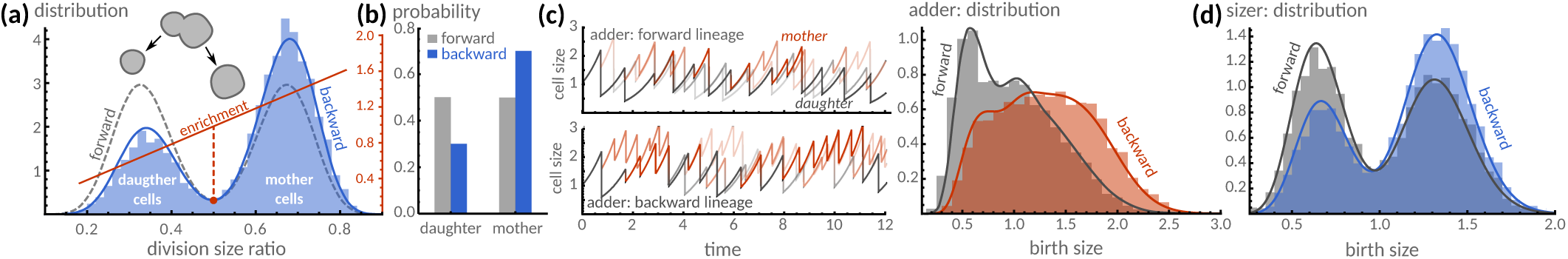
Asymmetric division increases the frequency of mother cells. A model population of budding yeast cells is described in which daughter cells inherit 1/3 of the mother size on average. **(a)** In forward lineages the relative division size indicates equal likelihoods for mother and daughter cells (dashed line). In contrast, our theory predicts that cells with inherited size fractions smaller than 1/2 (red dot) are under-represented while cells with larger ratios are over-represented in backward lineages (blue line). The enrichment (red line) represents the ratio of cells in backward and forward lineages, Eq. (15). **(b)** Cell size increases due to the increased likelihood of mother cells, which is well explained by their relative size proportions at cell division, Eq. (19). **(c)** A qualitative comparison of the adder mechanism in forward (top, left) and backward lineages (bottom, left) confirms that mother cells (red) are more frequent than daughter cells (black). This dependence gives rise to a broad distribution of birth sizes in backward lineages (right). **(d)** For the sizer control, asymmetric division leads to distinct sizes for mother and daughter cells that is qualitatively predicted by their inherited size-fraction (a). Simulations and theory assume Beta-distributed size-fractions with mean 1/3 and coefficient of variation of 20% for daughter cells with the rest attributed to mother cells.

Assuming size proportions of 1/3 for daughter cells and assigning the rest mother cells, we verify via stochastic simulations using the adder model that the proportion of daughter cells in backward lineage is indeed 1/3 (Fig. 3b). A qualitative comparison of simulated forward (Fig. 3c, top left) and backward lineages (c, bottom left) confirms this dependence, which gives rise to a broad distribution of birth sizes (c, right). For the sizer model (Fig. 3d) mother and daughter cells have well-separated size distributions, as expected.

### C. Populations of cells with varying cell-growth rates

We finally consider the case in which cells divide into daughters of equal sizes but differing cell-growth rates. The form of the division kernel describing this situation is

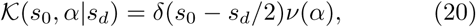

where *ν*(*α*) is the distribution of cell-growth rates in a forward lineage. In order to separate the contributions of size and cell-growth rate, we make the ansatz 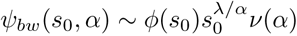, such that the backward lineage equation (9) simplifies to

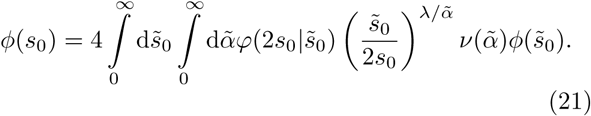

It is now clear that cell size in backward lineages depends on cell-growth rate. Integrating the above equation, one obtains an equation for λ that shows an intricate coupling between population growth, cell-growth rate and birth size that makes it difficult to analyse analytically.

#### Growth rate fluctuations are detrimental to population fitness

Although Eq. (21) needs to be solved numerically, upper and lower bounds of the population-growth rate can be obtained (Methods 3). Specifically, we find that the population-growth rate cannot exceed the mean growth rate of a single cell but the population doubling time is shorter than the time in which a single cell doubles its size, that is

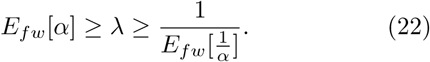

The underlying biological assumption for this result to hold is that cells cannot grow infinitely fast and it remains valid for asymmetrically dividing cell populations (Methods 3). Because the upper bound is sharp only for deterministic growth, it follows that fluctuations in cell-growth rate must decrease population growth.

It also follows that determining population-growth rate based on individual cell-growth rate always overestimate population-growth rate while measures based on size doubling rate always underestimate it. We validate this prediction using reported cell- and population-growth rates of *E. coli* cells^3,39^ (Fig. 4). Because these studies do not provide an estimate of 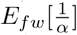, we used division time as a proxy (dashed red line). This yields the same lower bound as obtained by *Hashimoto et al.^3^.* Using simulations we found that the lower bound provided here, Eq. (22) can be sharper, especially for large division errors (not shown).

**FIG. 4.**
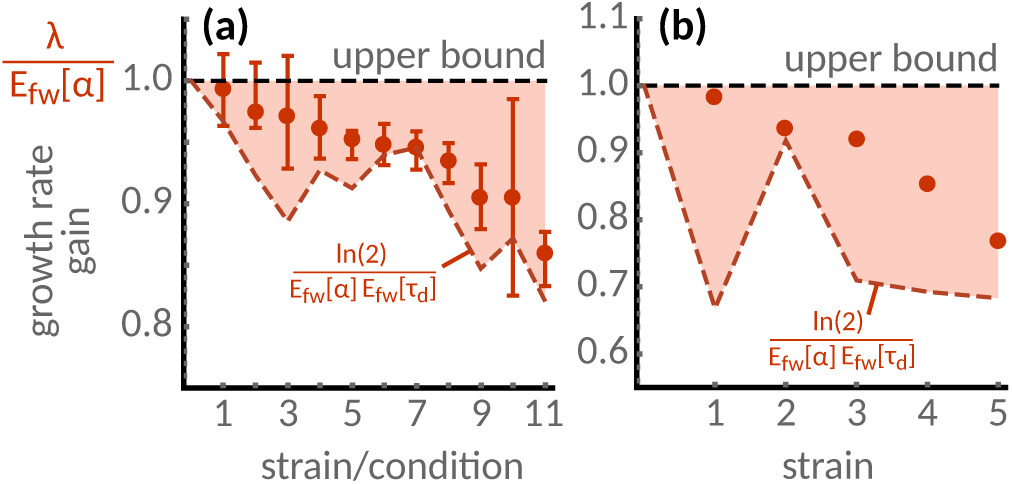
Measurements of *E. coli* cells validate limits on population growth. **(a)** Microfluidic experiments^3^ (red dots) confirm that the population-rate is smaller than the mean elongation rate of single cells (dashed black line). Strains and growth conditions are ranked by growth rate gain and error bars denote maximum error from the reported 95% bootstrap confidence intervals. Lower bounds are estimated using mean division times and elongation rates as in Ref.^3^ (dashed red line). **(b)** Time-lapse observations of microcolonies^39^ provide an independent experimental validation of the growth bounds.

To make further analytical progress, we ignore variability in cell size control, *φ*(*s_d_*|*s*_0_) ≈ *δ*(*s_d_* − 2*s*_0_), in the following. We then find that, after integrating, Eq. (21) reduces to

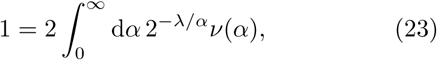

which determines the population growth rate λ. This equation is an Euler-Lotka equation^40^ with division time *τ_d_* = ln2/*α*. Its predictions are highly accurate for different modes of size control (Fig. 5a, solid line). Insights into its solution can be obtained by using an approximation valid for small cell-growth fluctuations (Methods 4), which results in

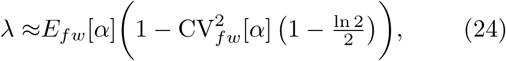

where *E_fw_*[*α*] is the expected growth rate of a single cell and CV_fw_[*α*] its coefficient of variation. Stochastic simulations verify the accuracy of this approximation in the physiological regime^4,5,26,39^ (CV_*fw*_ [*α*] < 0.5, Fig. 5a dot-dashed line).

To investigate the effect of cell size control on population growth, we numerically solve Eq. (21). We find that errors in size control can increase the population growth rate slightly above the estimate (23), which we demonstrate for the adder mechanism (Fig. 5a, inset). The impact of size control on population growth requires the presence of cell-growth variations, however. Thus the bounds given in Eq. (22) demonstrate that growth fluctuations fundamentally constrain population fitness.

**FIG. 5.**
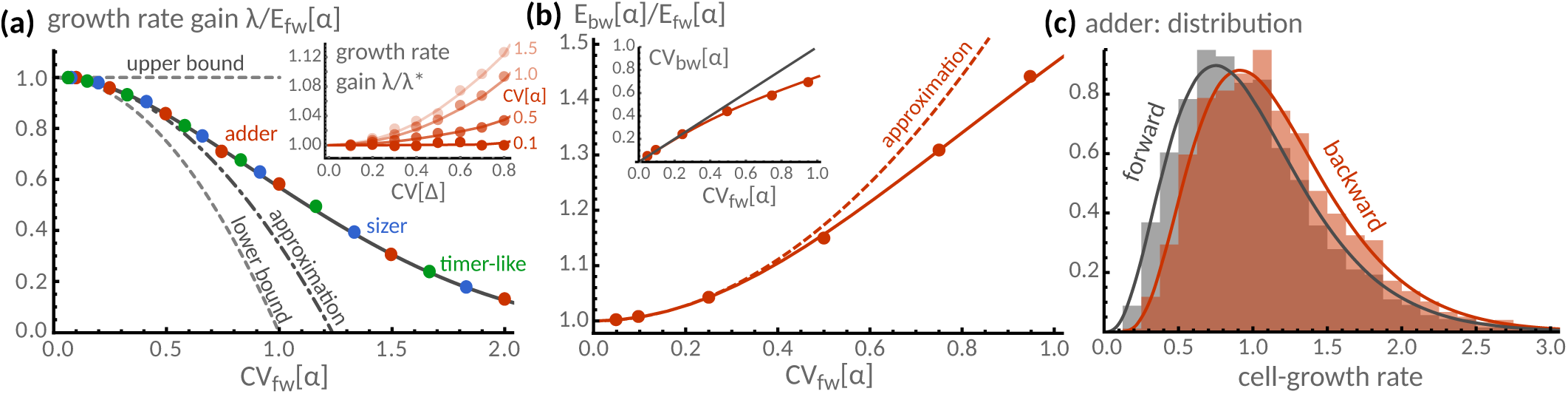
Cell-growth rate fluctuations affect population fitness. **(a)** Gain in population-growth rate λ/*E_fw_* [*α*] obtained from stochastic simulations of sizer (blue *a* = 0), adder (red *a* = 1), and timer-like size controls (green dots *a* = 1.5). For small errors in the size control, all of these data collapse onto a single line (solid black) given by the solution of Eq. (23). The approximation valid in the limit of small fluctuations, Eq. (24), captures the behaviour in the physiological regime (0 − 0.5). For comparison, we show theoretical bounds (dashed grey). (inset) Variations in size control, shown for the adder model with increments Δ, can slightly increase population growth above the estimate λ* of Eq. (23), but require the presence of cell-growth rate fluctuations. **(b)** The expected cell-growth rate in backward lineages increases with the size of growth rate fluctuations relative to forward lineages. The solutions to Eq. (25) (solid red) is verified against simulations using an adder mechanism (dots) and an approximation (dashed red), Eq. (26). The size of fluctuations in these lineages is generally smaller than expected for single cells (inset). **(c)** Lineage distributions are compared for both lineage types with a coefficient of variation of 0.5. Theory (lines) and simulations (dots, shaded areas) assume Gamma-distributed cell-growth rates with unit mean and prescribed coefficient of variation; size control errors are modelled using a Gamma distribution with coefficient of variation of 10% except in (a, inset).

#### Cells grow faster in backward lineages

Having determined the population growth rate, we discuss the distribution of cell-growth rates in a backward lineage. Since this distribution determines the cell-growth rate, it must be given by the term below the integral of Eq. (23),

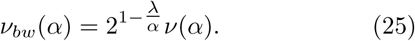

Interestingly, we find that the probability of cell-growth rates in forward and backward lineages is equal for cells growing at the same rate as the population. As a consequence cells growing faster than the population are over-represented while slower cells are under-represented. Using a similar approximation as in Eq. (24), we find that

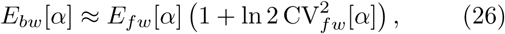

meaning that the expected cell-growth rate in a backward lineage is larger than in a forward lineage (Fig. 5b dashed red line). The resulting shift to higher cell-growth rates in the backward lineage distributions (Fig. 5c) is well described by Eq. (25). Similarly, the size of fluctuations is smaller in backward than in forward lineages because of the increased mean (Fig. 5b, inset). Our predictions are shown to be in excellent agreement with stochastic simulations.

**FIG. 6.**
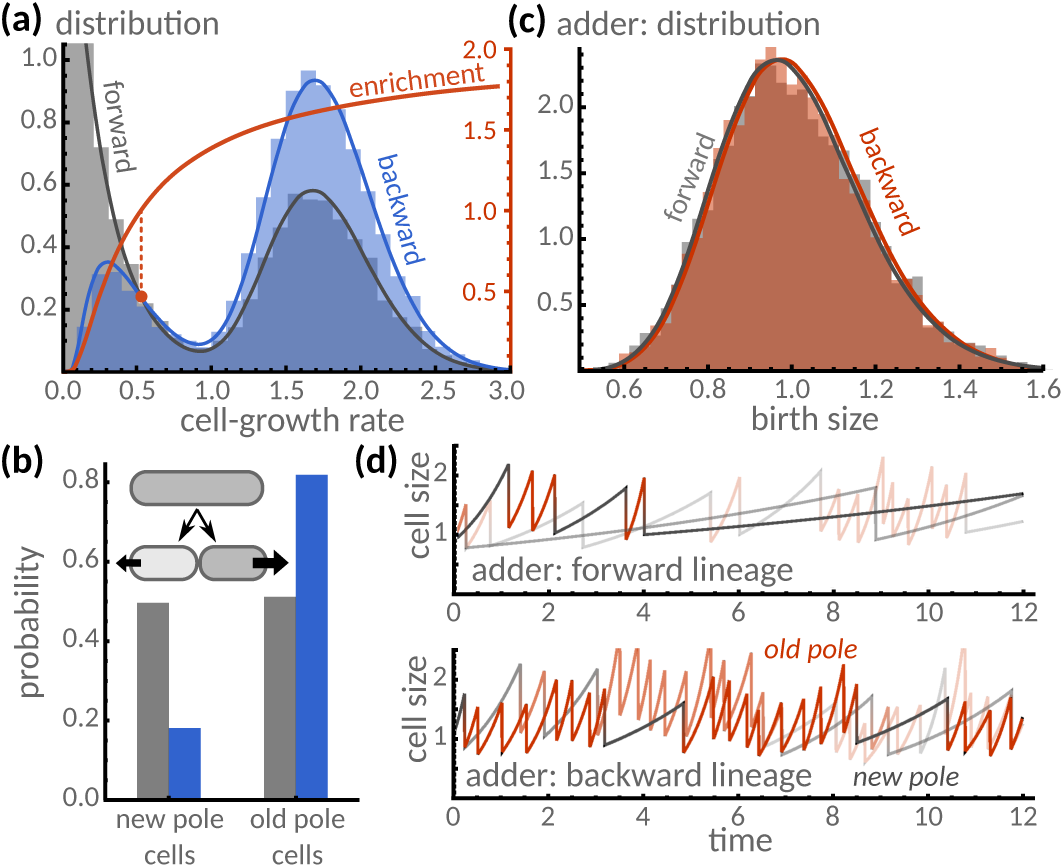
Asymmetric growth shifts the frequency of growth phenotypes. A situation is described in which old pole cells grow faster than new pole cells. **(a)** Asymmetric growth tilts the distribution of cell-growth rates in forward lineages (black line) towards fast growing cells in backward lineages (blue line). Specifically, cells with growth-rates above the population-growth rate (red dot) are over-represented while cells below are under-represented. This enrichment (red line) is in excellent agreement with the theoretical predictions (Eq. (25)). **(b)** Proportions of new and old pole cells show less slow growing cells than fast growing cells. **(c)** Birth-size distributions of both lineage types agree and resemble ideal lineages. **(d)** Simulated lineages qualitatively confirm the enrichment of old (red) compared to new pole cells (black) in backward lineages. Theory and simulations assume Gamma-distributed adder control with a coefficient of variation of 30%.

#### Histories of asymmetrically growing cells contain fewer slow than fast growing cells

Single-cell experiments revealed that asymmetric division in mycobacteria results in sister cells that differ in their elongation rates^41,42^. Specifically, a daughter inheriting the new pole grows slower than an old pole cell. By symmetry it is clear that the proportions of old pole (op) and new pole cells (np) must be equal in forward lineages,

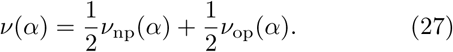

The distribution of individual cell-growth rates in backward lineages is obtained by using the above relation together with Eq. (25). For new pole and old pole cells with different mean growth rates, Fig. 6a (left) shows that the growth rate distribution in backward lineages contains fewer cells with a slow growing pole than expected in forward lineages. The fraction of new pole cells observed in backward lineages is given by an exponential weighting of the cell-growth rate against the population-growth rate

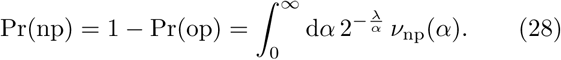

To understand this result in more detail, define the size doubling time *τ_d_* = ln2/*α* and note that 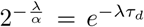 is the size-proportion that a cell growing at rate *α* contributes to the growth of the population. We verify this dependence through simulation (Fig. 6b). We also show that the distribution of birth sizes is indeed close to the statistics of ideal lineages (c). The good agreement with the simulations verifies the accuracy of the employed approximations. Qualitatively, a few stochastic realisations of forward (d, top) and backward lineages (bottom) confirm this result.

## III. DISCUSSION

We presented a method to predict histories of single cells in an exponentially growing population. Our analysis reveals that physiological differences in sister cells have a significant impact on individual cell histories and their contribution to the overall population-growth. We found that if every cell division results in a pair of perfectly identical sister cells, inheriting both the growth rate and equal proportions of the mother’s size, the statistics of forward lineages histories can be predictive for cell histories. Cell division is naturally imperfect and when this happens, cell histories in a population deviate from forward lineages because larger cells grow faster and hence contribute more to the overall population growth, which becomes particularly conspicuous for asymmetrically dividing cells such as budding yeast. When sister cells grow at different rates we showed that it is the size-fraction that a cell contributes to the growth of the population that determines its history. Consequently, fast growing cells contribute more to a typical cell’s history than they contribute population growth. These findings highlight the dependence of cellular histories on physiolgy.

A range of single-cell tracking devices are used to study cellular growth and size control. Mother machines, for instance, are used to measure forward lineages of *E. coli* cells. Such devices yield qualitatively different results from chemostats, as for example, increased filamentation rates^43^ and the production of mini-cells^9^. Our theory could be used to explain these differences because of the different proportions cells are being represented in cell histories and growing populations. Microchemostats^3,14,43^ are ideal candidates to facilitate the long-term observation of cell histories that we have proposed here. More recent approaches allow measuring growth rates directly in populations, the statistics of which can be inferred from cellular histories (Methods 2). The proposed quantitative framework thus bridges the gap between these competing experimental approaches^44^.

Using this theory, we obtained two fundamental bounds on the population dynamics: the population growth-rate can never exceed the rate at which an average cell grows but the population-doubling time is shorter than the time at which an average cell doubles its size. We validated this dependence using available single-cell data. Within the theoretical bounds, cell size control can affect population growth but only in the presence of cell-growth variations. It hence follows that growth fluctuations are fundamentally limiting the fitness of a cell population.

Previous studies provided similar lower bounds on population-growth rate based on division timing^3,40^. We found that when cells control their division size, leading to correlated division times, that cell populations cannot exploit their phenotypic variations to increase population fitness above the rate at which an individual cell grows, which we validated using available single-cell data. Populations could exploit this mechanism for noise reduction by negative selection, for instance, through mutations that tend to decrease individual growth differences.

Individual cells must make decisions based on their past. The fact that cells within a clonal population experience histories that cannot be predicted solely from observations of cells in isolation is due to the division process that results in progeny with variable physiology. It would be interesting to study how cells may exploit these historical fluctuations to win competitions or to enable population-level strategies^45,46^.

## ACKNOWLEDGEMENTS

It is a pleasure to thank Andrea Y. Weiße and Vahid Shahrezaei for stimulating discussions. PT gratefully acknowledges support by The Royal Commission for the Exhibition of 1851.

## METHODS

### 1. Statistics of forward lineages

We first summarize the lineage approach briefly. We consider a sequence of cellular birth sizes where the information about one of the daughters is discarded. The sequence of birth volumes and cell-growth rates obtained in this way is a forward lineage. In simple terms, it can be modelled by expressing the birth size at the start of the (*n* + 1)-th cell cycle as a function of the size at the end of the *n*-th cell cycle^16,19^

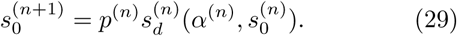

We here assume that (i) the proportions of size *p*^(*n*)^ inherited by a daughter cell of the *n*-th is identically and independent distributed with density *π*(*p*), (ii) the cell-growth rates *α*^(*n*)^ are independently distributed between daughter cells with distribution *ν*(*α*), and (iii)the division sizes 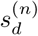 are conditionally independent with density *φ*(*s_d_*|*s*_0_,*α*). The stationary lineage distribution *ψfw*(*s*_0_, *α*) is obtained by writing an integral equation that maps the *n*-th generation’s distribution to the (*n* + 1)-th generation’s distribution and taking the limit *n* → ∞. The result is Eq. (4).

#### Characterising cell size control using linear models

On a phenomenological level, we assume that division size is independent of cell-growth rate and set *φ*(*s_d_*|*s*_0_,*α*) = *φ* (*s_d_* − *as*_0_). This corresponds to a linear model^11,25^ of the function 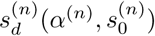 in Eq. (29) given by

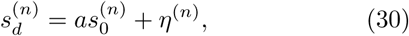

where the parameter *a* implements the size control (*a* = 0 sizer, *a* =1 adder, *a* = 2 timer, see Sec. IIB) and the *η*^(*n*)^ are independent and identically distributed increments.

For any cell in a forward lineage, the distribution of growth rate is then 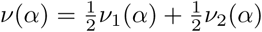 but the lineage distribution of birth sizes is difficult to obtain analytically. It is often more straightforward to obtain a characterisation in terms of moments^17^. The mean birth size can be obtained by using the linear model in Eq. (4) and multiplying the result by *s*_0_ and performing the integration. This shows that

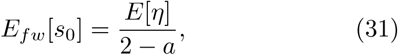

is independent of the division error CV[*p*]. However, the mean birth size exists only if *a* < 2, and thus ruling out a timer mechanism for size-homoeostasis^19,47,48^.

### 2. Statistics of backward lineages

The density *n*(*τ, s,α,t*)d*τ*d*s*d*α* counts the number of cells at time *t* with age between *τ* and *τ*+d*τ*, size between *s* and *s*+d*s*, and cell-growth rate between *α* and *α*+d*α*. Denoting by *γ*(*τ, s, α*)d*τ*d*s*d*α* the probability for a cell to divide at age *τ* and size *s*, the cell density evolves as follows

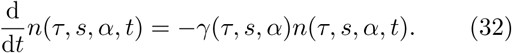

When the cell-growth rate *α* does not change between cell divisions, expanding the total time derivative in Eq. (32) leads to the evolution equation (7a) given in the main text. The boundary condition, Eq. (7b), ensures that the number of newborn cells must equal the number of dividing cells.

Integrating Eq. (32) shows that the distribution of division times *τ*_d_ is given by Eq. (2) of the main text. To characterise the single cell behaviour, we change from division times to division sizes characterised by the distribution *φ*(*s*|*s*_0_,*α*) given by Eq. (3). While this characterisation is exactly equivalent to Eq. (2) (see also Ref.^29^), it allows for a tremendous simplification of Eqs. (7) because it removes the simultaneous depends on cell age and size. The result of this change of variables is the evolution equation

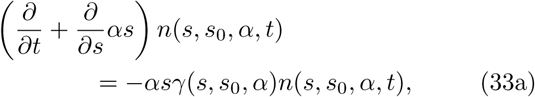

which is subject to the boundary condition

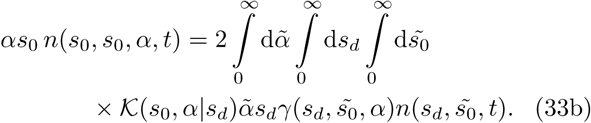

The solution of Eqs. (33) can in principle be obtained by separating variables. Formally, we write

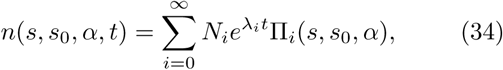

where λ_*i*_ denote the eigenvalues of the integro-differential operator. In the long-term limit the solution is dominated by the eigenvalue with largest real part λ_0_. For now, we assume that such a dominant eigenvalue exists and set

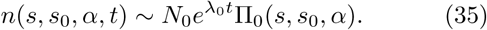

We identify Π_0_(*s, s*_0_, *α*) as the stable distribution of size, birth size and cell-growth rate. We then see from Eq. (6) of the main text that the total cell number grows exponentially *N*(*t*) ~ *N*_0_*e*^λ_0_*t*^ for long times. Note that in the main text we use λ to denote the dominant eigenvalue instead of λ_0_. Using the above equation in Eq. (33a), we find

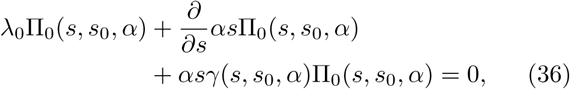

which can be solved straightforwardly

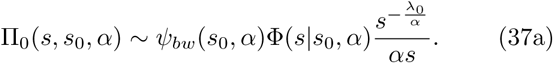

Here Φ(*s, s*_0_, *α*) is related to the cumulative distribution function of *φ* via

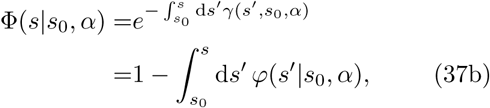

which gives the probability that a cell with birth size *s*_0_ and growth rate *α* has not divided before reaching size *s*. The density *ψ*_*bw*_ (*s*_0_, *α*) is the ancestral distribution of cell-growth rate and birth size that contains the information about the history of the cells. *ψ*_*bw*_(*s*_0_, *α*) is to be determined from the boundary conditions and the denominator containing *α* is a factor chosen for convenience. Inserting Eq. (35) with Π_0_(*s, s*_0_, *α*) into Eq. (33b), we find

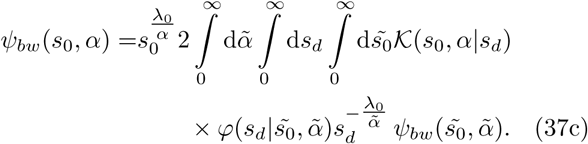

This equation determines (i) the population-growth rate λ_0_ and (ii) the distribution of birth sizes in a backward lineage.

### 3. Bounds on the population-growth rate

To obtain lower and upper bounds on the population-growth rate we specialise Eq. (37c) to the case in which growth rate fluctuations and division errors are independent

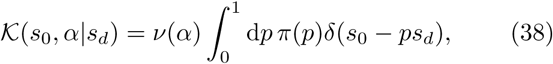

where *ν*(*α*) is the distribution of growth rates and *π*(*p*) is the ratio of daughter to mother size after cell division in a forward lineage. As in the main text, we now set 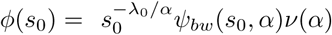 and assume that 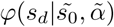 to be independent of 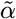. We then obtain

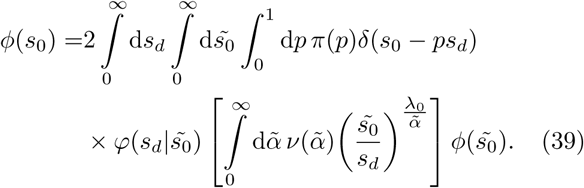

Computation of the bounds requires bounding the expectation value in square brackets.

#### a. Lower bound

We notice that the integrand of the 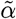-integral is convex in 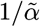. Using Jensen’s inequality we obtain

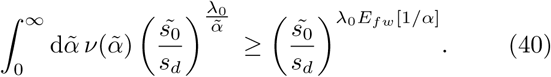

Inserting the above relation into Eq. (39) we obtain the inequality

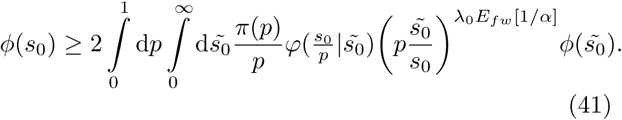

Multiplying this inequality by 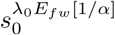 and integrating over *s*_0_ we find

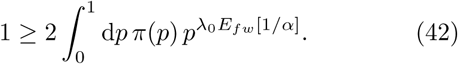

Note that because *π*(*p*) = π(1 − *p*) equality holds when λ_0_ equals 1/*E_fw_* [1/α]. Larger values of λ_0_ decrease the integral and hence

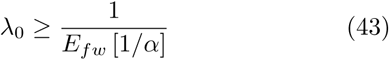

is a lower bound.

#### b. Upper bound

To obtain an upper bound we set 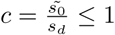 and observe that

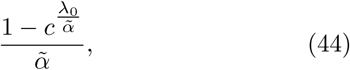

is a convex function for λ_0_ > 0 and 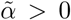. Restricting ourselves to distributions with finite means and variances, we can employ bounds derived in Ref. ^49^ leading to

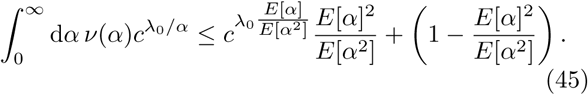

The distribution that uniquely achieves this bound is

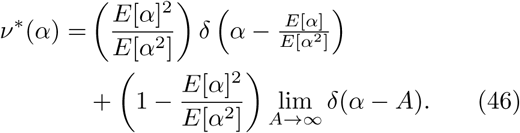

We are now faced with two options: either we allow for distributions with a finite probability mass at infinity, or the only admissible distribution is the deterministic one, for which *E*[*α*^2^] = *E*[*α*^2^] and the second term in the above equation vanishes. Certainly, the latter situation is biologically relevant since single-cell growth rates are bounded^50,51^. It then follows by using Eq. (45) in (39) that the deterministic distribution bounds the population growth rate from above and thus

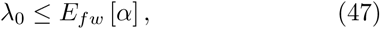

following the same arguments as in Sec. II B.

### 4. Perturbative calculation of population growth rate

Next, we derive an approximation for the eigenvalues determined by the characteristic equation (23). We set *α*= *α*_0_+*σξ*, where *α*_0_ is the average cell-growth rate, its standard deviation is *σ* and *ξ* denotes a random variable of zero mean and unit variance. Inserting λ = *α*_0_+*σ*λ_1_+*σ*^2^λ_2_+*Ο*(*σ*^3^) into Eq. (23) and truncating of terms of order *σ*^0^ we find

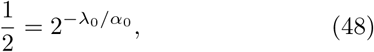

whose solutions are 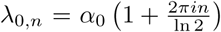 for *n* = 0; 1;…Thus to this order there exists no eigenvalue with dominant real part. Truncation after terms of order *σ* yields λ_1_,_*n*_ = 0 because *E*[*ξ*] = 0. Truncating the characteristic equation after terms of order *σ*^2^, we find

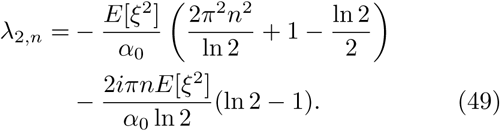

Because the real part of Eq. (49) decreases with *n* we find that λ_0_ is the dominant eigenvalue. Letting *n* = 0 in Eq. (49) we obtain the approximation given in Eq. (24) of the main text by expressing the result in terms of the mean, and the coefficient of variation of *α*.

## References

1 F. Ferrezuelo, N. Colomina, A. Palmisano, E. Garí, C. Gallego, A. Csikász-Nagy, and M. Aldea, “The critical size is set at a single-cell level by growth rate to attain homeostasis and adaptation,” Nat Commun 3, 1012 (2012).

2 S. Taheri-Araghi, S. Bradde, J. T. Sauls, N. S. Hill, P. A. Levin, J. Paulsson, M. Vergassola, and S. Jun, “Cell-size control and homeostasis in bacteria,” Curr Biol 25, 385–391 (2015).

3 M. Hashimoto, T. Nozoe, H. Nakaoka, R. Okura, S. Akiyoshi, K. Kaneko, E. Kussell, and Y. Wakamoto, “Noise-driven growth rate gain in clonal cellular populations,” Proc Natl Acad Sci, 201519412 (2016).

4 M. Wallden, D. Fange, E. G. Lundius, Ö. Baltekin, and J. Elf, “The synchronization of replication and division cycles in individual E. coli cells,” Cell 166, 729–739 (2016).

5 D. J. Kiviet, P. Nghe, N. Walker, S. Boulineau, V. Sunderlikova, and S. J. Tans, “Stochasticity of metabolism and growth at the single-cell level,” Nature 514, 376–379 (2014).

6 V. Shahrezaei and S. Marguerat, “Connecting growth with gene expression: of noise and numbers,” Curr Opin Microbiol 25, 127–135 (2015).

7 J. M. Raser and E. K. O’Shea, “Noise in gene expression: origins, consequences, and control,” Science 309, 2010–2013 (2005).

8 G. Lambert and E. Kussell, “Quantifying selective pressures driving bacterial evolution using lineage analysis,” Phys Rev X 5, 011016 (2015).

9 P. Wang, L. Robert, J. Pelletier, W. L. Dang, F. Taddei, A. Wright, and S. Jun, “Robust growth of Escherichia coli,” Curr Biol 20, 1099–1103 (2010).

10 M. M. Crane, I. B. Clark, E. Bakker, S. Smith, and P. S. Swain, “A microfluidic system for studying ageing and dynamic singlecell responses in budding yeast,” PloS one 9, e100042 (2014).

11 Y. Tanouchi, A. Pai, H. Park, S. Huang, R. Stamatov, N. E. Buchler, and L. You, “A noisy linear map underlies oscillations in cell size and gene expression in bacteria,” Nature 523, 357–360 (2015).

12 I. Soifer, L. Robert, and A. Amir, “Single-cell analysis of growth in budding yeast and bacteria reveals a common size regulation strategy,” Curr Biol 26, 356–361 (2016).

13 E. J. Stewart, R. Madden, G. Paul, and F. Taddei, “Aging and death in an organism that reproduces by morphologically symmetric division,” PLoS Biol 3, e45 (2005).

14 G. Ullman, M. Wallden, E. G. Marklund, A. Mahmutovic, I. Razinkov, and J. Elf, “High-throughput gene expression analysis at the level of single proteins using a microfluidic turbidostat and automated cell tracking,” Philos Trans R Soc 368, 20120025 (2013).

15 D. M. Wolf, L. Fontaine-Bodin, I. Bischofs, G. Price, J. Keasling, and A. P. Arkin, “Memory in microbes: quantifying history-dependent behavior in a bacterium,” PLOS one 3, e1700 (2008).

16 N. Brenner and Y. Shokef, “Nonequilibrium statistical mechanics of dividing cell populations,” Phys Rev Lett 99, 138102 (2007).

17 A. Amir, “Cell size regulation in bacteria,” Phys Rev Lett 112, 208102 (2014).

18 A. Marantan and A. Amir, “Stochastic modeling of cell growth with symmetric or asymmetric division,” arXiv preprint arXiv:1602.01848 (2016).

19 C. A. Vargas-Garcia and A. Singh, “Hybrid systems approach to modeling stochastic dynamics of cell size,” bioRxiv, 044131 (2016).

20 B. Charlesworth et al., Evolution in age-structured populations, Vol. 2 (Cambridge University Press Cambridge, 1994).

21 J. Hein, M. Schierup, and C. Wiuf, Gene genealogies, variation and evolution: a primer in coalescent theory (Oxford University Press, USA, 2004).

22 S. Leibler and E. Kussell, “Individual histories and selection in heterogeneous populations,” Proc Natl Acad Sci 107, 13183–13188 (2010).

23 Y. Wakamoto, A. Y. Grosberg, and E. Kussell, “Optimal lineage principle for age-structured populations,” Evolution 66, 115–134 (2012).

24 M. Campos, I. V. Surovtsev, S. Kato, A. Paintdakhi, B. Beltran, S. E. Ebmeier, and C. Jacobs-Wagner, “A constant size extension drives bacterial cell size homeostasis,” Cell 159, 1433–1446 (2014).

25 J. T. Sauls, D. Li, and S. Jun, “Adder and a coarse-grained approach to cell size homeostasis in bacteria,” Curr Opin Cell Biol 38, 38–44 (2016).

26 S. Tsuru, J. Ichinose, A. Kashiwagi, B.-W. Ying, K. Kaneko, and T. Yomo, “Noisy cell growth rate leads to fluctuating protein concentration in bacteria,” Phys Biol 6, 036015 (2009).

27 M. Osella, E. Nugent, and M. C. Lagomarsino, “Concerted control of Escherichia coli cell division,” Proc Natl Acad Sci 111, 3431–3435 (2014).

28 G. I. Bell and E. C. Anderson, “Cell growth and division: I. A mathematical model with applications to cell volume distributions in mammalian suspension cultures,” Biophys J 7, 329 (1967).

29 O. Diekmann, H. Lauwerier, T. Aldenberg, and J. Metz, “Growth, fission and the stable size distribution,” J Math Biol 18, 135–148 (1983).

30 M. Rading, T. Engel, R. Lipowsky, and A. Valleriani, “Stationary size distributions of growing cells with binary and multiple cell division,” J Stat Phys 145, 1–22 (2011).

31 E. B. Stukalin, I. Aifuwa, J. S. Kim, D. Wirtz, and S. X. Sun, “Age-dependent stochastic models for understanding population fluctuations in continuously cultured cells,” J R Soc Interface 10, 20130325 (2013).

32 D. Ramkrishna and M. R. Singh, “Population balance modeling: Current status and future prospects,” Annu Rev Chem Biomol Eng 5, 123–146 (2014).

33 L. Robert, M. Hoffmann, N. Krell, S. Aymerich, J. Robert, and M. Doumic, “Division in Escherichia coli is triggered by a size-sensing rather than a timing mechanism,” BMC Biol 12, 1 (2014).

34 J. A. Metz and O. Diekmann, The dynamics of physiologically structured populations, Vol. 68 (Springer, 1986).

35 H. J. Heijmans, “On the stable size distribution of populations reproducing by fission into two unequal parts,” Math Biosci 72, 19–50 (1984).

36 L. Koppes, C. L. Woldringh, and N. Nanninga, “Size variations and correlation of different cell cycle events in slow-growing Escherichia coli.” J Bacteriol 134, 423–433 (1978).

37 F. J. Trueba, “On the precision and accuracy achieved by Escherichia coli cells at fission about their middle,” Arch Microbiol 131, 55–59 (1982).

38 J. M. Guberman, A. Fay, J. Dworkin, N. S. Wingreen, and Z. Gitai, “Psicic: noise and asymmetry in bacterial division revealed by computational image analysis at sub-pixel resolution,” PLoS Comput Biol 4, e1000233 (2008).

39 A. S. Kennard, M. Osella, A. Javer, J. Grilli, P. Nghe, S. J. Tans, P. Cicuta, and M. C. Lagomarsino, “Individuality and universality in the growth-division laws of single e. coli cells,” Phys Rev E 93, 012408 (2016).

40 P. Painter and A. Marr, “Mathematics of microbial populations,” Annu Rev Microbiol 22, 519–548 (1968).

41 B. B. Aldridge, M. Fernandez-Suarez, D. Heller, V. Ambravaneswaran, D. Irimia, M. Toner, and S. M. Fortune, “Asymmetry and aging of mycobacterial cells lead to variable growth and antibiotic susceptibility,” Science 335, 100–104 (2012).

42 K. J. Kieser and E. J. Rubin, “How sisters grow apart: mycobacterial growth and division,” Nat Rev Microbiol 12, 550–562 (2014).

43 Z. Long, A. Olliver, E. Brambilla, B. Sclavi, M. C. Lagomarsino, and K. D. Dorfman, “Measuring bacterial adaptation dynamics at the single-cell level using a microfluidic chemostat and timelapse fluorescence microscopy,” Analyst 139, 5254–5262 (2014).

44 N. Cermak, S. Olcum, F. F. Delgado, S. C. Wasserman, K. R. Payer, M. A. Murakami, S. M. Knudsen, R. J. Kimmerling, M. M. Stevens, Y. Kikuchi, et al., “High-throughput measurement of single-cell growth rates using serial microfluidic mass sensor arrays,” Nat Biotechnol (2016).

45 J.-W. Veening, E. J. Stewart, T. W. Berngruber, F. Taddei, O. P. Kuipers, and L. W. Hamoen, “Bet-hedging and epigenetic inheritance in bacterial cell development,” Proc Natl Acad Sci 105, 4393–4398 (2008).

46 B. M. Martins and J. C. Locke, “Microbial individuality: how single-cell heterogeneity enables population level strategies,” Current opinion in microbiology 24, 104–112 (2015).

47 G. I. Bell, “Cell growth and division: III. Conditions for balanced exponential growth in a mathematical model,” Biophys J 8, 431 (1968).

48 K. B. Hannsgen, J. J. Tyson, and L. T. Watson, “Steady-state size distributions in probabilistic models of the cell division cycle,” SIAM J Appl Math 45, 523–540 (1985).

49 B. Guljaš, C. E. Pearce, and J. Pečarić, “Jensen’s inequality for distributions possessing higher moments, with application to sharp bounds for Laplace-Stieltjes transforms,” J Austral Math Soc B 40, 80–85 (1998).

50 M. Scott, C. W. Gunderson, E. M. Mateescu, Z. Zhang, and T. Hwa, “Interdependence of cell growth and gene expression: origins and consequences,” Science 330, 1099–1102 (2010).

51 A. Y. Weiße, D. A. Oyarzún, V. Danos, and P. S. Swain, “Mechanistic links between cellular trade-offs, gene expression, and growth,” Proc Natl Acad Sci 112, E1038–E1047 (2015).

